# Inventory of cultivatable populations of S-cycling, fermentative, Fe-reducing, and aerobic heterotrophic bacteria from salt marsh sediments

**DOI:** 10.1101/048611

**Authors:** David Emerson, Cynthia M. Lydell

## Abstract

A survey was carried out of the dominant chemotrophic groups of bacteria inhabiting surface salt marsh sediments in the Virginia Coastal Reserve (VCR) on the Atlantic coast of Virginia. Total direct cell counts were carried out on all samples. Aerobic heterotrophs, sulfur oxidizers, sulfate reducing bacteria (SRB), sulfur disproportionaters, Fe-reducing (FeRB) and fermentative bacteria were all quantified by most probable number (MPN) at four different sites that ranged in spatial scale from a few meters to 15 km apart. The sites were sampled every 3 – 4 months over a two year period. Total cell counts were quite consistent temporally at each of the sites, and ranged from a high of 1.4 × 10^10^ cells. gdw^-1^ to a low of 8 × 10^8^ cells. gdw^-1^. Recoveries of all culturable bacteria were also site dependent and ranged from a minimum of 0.4% to a maximum of 40% of the total cell count. Aerobic bacteria were the dominant recovered population at all of the sites, followed by sulfur-oxidizing bacteria. Together these two groups accounted for >75% of the total recovered bacteria at each of the sites. The populations of anaerobic groups fluctuated significantly; S-disproportionating and SRB were most abundant followed by FeRB and fermenters. On average, all the anaerobes were in the same order of magnitude of abundance (10^7^ cells. gdw^-1^). Overall, these results suggest that aerobic bacteria consistently predominated in the top 10 cm of the marsh sediments, and that autotrophy related to sulfur oxidation and disproportionation may be important, but under studied processes in salt marsh ecosystems.

## INTRODUCTION

Coastal salt marshes can be among the most productive of ecosystems in terms of the amount of C-produced per m^2^. While their precise role in the nutrient dynamics of associated larger coastal ecosystems remains a source of debate(Odum 2000), it is agreed that salt marshes are important nurseries for the larval and juvenile stages of a number of commercially important fin and shellfish species (Deegan 2000). Furthermore salt marshes are important physical buffers that can act to ameliorate storm waves and tidal surges thus protecting the fragile inshore coast line. The macrofauna and microfauna that comprise the marsh and its sediments play an important role in maintaining this physical integrity. The United States Environmental Protection Agency has reported that one consequence of global warming will be radical alteration of coastal marshes due to sea level rise along the Eastern Seaboard of the United States (Anonymous 2002), lending a new imperative to the study of these resilient, yet potentially endangered ecosystems.

Despite a monotonous plant cover dominated by *Spartina alterniflora*, salt marshes are home to a diverse and abundant community of microorganisms that mineralize plant matter using a variety of electron acceptors and pathways. The most intensively studied group of salt marsh microbes are the sulfate reducing bacteria (SRB) (Rooney-Varga et al. 1998, Hines et al. 1999, Kostka et al. 2002a, Purdy et al. 2002). More recently FeRB have also received attention (Lowe et al. 2000, Kostka et al. 2002b, Koretsky et al. 2003). Lovell and co-workers have carried out detailed molecular and cultural studies on diazotrophic bacteria in salt marsh sediments in South Carolina (Bagwell et al. 2000, Piceno et al. 2000a, b, Lovell et al. 2001). Recently these same workers showed that diverse functional sequences for the formyltetrahydrofolate synthase genes indicative of acetogenic bacteria were ubiquitous in the rhizosphere of salt marsh plants (Leaphart et al 2003). Another recent study used fluorescence in situ hybridization (FISH) to quantitate different microbial groups in rhizosphere sediments associated with a mesohaline Spartina salt marsh and an immediately adjacent region of marsh in which Phragmites was dominant (Burke et al 2002). In general, these latter studies indicate that populations of salt marsh microbes while diverse appear quite stable throughout the seasons. However, there have been surprisingly few attempts at quantitating other potentially important microbial groups in salt marsh sediments, including S-oxidizing and S-disproportionating bacteria, aerobic heterotrophs, and fermentative bacteria.

As part of a biotic survey of microbes in Atlantic coastal salt marshes, we have systematically attempted to assess the culturable populations of dominant chemotrophic microbes from salt marsh sediments both temporally and spatially. In the marshes we investigated, our general findings are consistent with the above mentioned theme of relatively consistent population sizes and structures. Furthermore, population sizes for culturable members of under studied but important microbial groups including S-oxidizing bacteria, S-disproportionating bacteria, fermentative bacteria, and, aerobic heterotrophs are reported, as well as numbers for SRB, and FeRB. These data provide an important baseline for both microbial community studies and biogeochemical studies in salt marsh ecosystems.

### METHODS AND MATERIALS

#### Sites

The VCR is a long term ecological research site maintained by the University of Virginia. Four sites that were separated on a scale of meters to kilometers were chosen for repeated sampling and are shown in Fig 1. These sites were marked and at each sampling time cores were taken within 3 m of the marker. Physical characteristics of these sites are given in Table 1. The Hog Island S1 site was in a tall form *Spartina* marsh a few meters from the upland margin on Hog Island. The Hog Island S2 site was about 4 meters from the S1 site, located in a small pool (approx. 2 × 4 m) that was devoid of plant growth, although the sediment did contain roots from the surrounding *Spartina*. The sediment at this site was a thick black mud that often smelled of sulfide; in the winter white precipitates of elemental S were visible on the surface of the mud. The third site, Cattleshed, was located in a large open marsh approximately 1.5 km from the Hog Island sites. It was dominated by short form *Spartina*, and the sediment was a homogeneous gray mud. The fourth site, Red Bank, was on a creek bank adjacent to the mainland about 15 km from Hog Island. The sediment at this site was also gray colored and appeared to have a very high clay content. The site was at the edge of a growth of *Spartina*; immediately above this the vegetation shifted to *Distyclis*.

**Table 1.** Site Characteristics.

| Site | Location | %Organic Matter | CHN | Temp. Range | Sediment pH | Salinity | Color/Texture |
| --- | --- | --- | --- | --- | --- | --- | --- |
| Hog Is, S1 | 37° 27.167' N<br>75° 40.513' W | 3.96 | 1.32: 0.13:<br>0.09 | 8 - 29 | 7.03 | ND<sup>a</sup> | Gray - brown,<br>clay/sand |
| Hog Is, S2 | 37° 27.167' N<br>75° 40.513' W | 44.0 | 9.38: 1.31:<br>0.70 | 5 - 30.5 | 7.28 | 30 | Black, sand/clay |
| Cattleshed | 37° 26.595' N<br>75° 41.315' W | 5.42 | 1.73: 0.33:<br>0.19 | 13 - 27 | 7.01 | 33 | Gray, clay/sand |
| Red Bank | 37° 26.731' N<br>75° 50.382' W | 3.73 | 0.85: 0.21:<br>0.09 | 14 - 30 | 6.99 | 35 | Gray, clay |

<sup>a</sup>ND, not determined.

a ND, not determined.

**Fig. 1.**
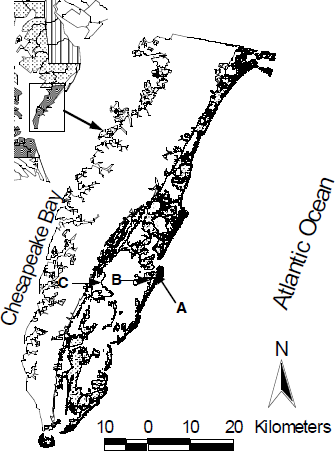
Map of the Virginia Coastal Reserve. Sampling sites are marked with arrows, A is the Hog Island Sites 1 and 2; B is the Cattleshed site, and C is the Red Bank site.

#### Sampling

Samples were collected using sterilized core barrels constructed from 3 inch diameter PVC pipe that was sharpened at one end. The cores were pushed by hand approximately 15 cm into the sediment, and intact sediment cores were extracted. The cores were stored on ice. All processing of the cores was done in an anaerobic glove box (Coy Laboratory Products, Grass Lake, MI) containing 97% N_2_ : 3% H_2_ within 12-24 h of collection. The top 5 – 7 cm of the sediment was removed and enough filtered and autoclaved seawater (collected from the site) was added to yield a slurry that could be pipeted easily. The slurry was used to set-up dilution series for inoculation of most probable number (MPN) tubes or for quantitative plating. All samples were normalized to a gram dry wt (gdw) basis by drying measured amounts of the slurry (in triplicate) at 85°C for at least 72h. Sediment organic matter content was calculated by loss on ignition at 430°C for 16 hours (Nelson and Sommers 1982). Total sediment C and N were determined using a CHN analyzer (Perkin Elmer Series II 2400). The pH at the sampling sites was determined in situ with a hand held pH meter (Cole-Parmer pH/CON10); the probe was placed in the core hole immediately after removal of the core. Salinity was determined on site with a hand held refractometer.

#### Quantitative culture work

Eight different metabolic groups of organisms were enumerated using 3 tube or 5 tube MPNs (de Man 1975). All MPN dilutions were done using Eppendorf pipettors with barrier pipet tips to reduce the risk of contamination. All MPNs were done in glass tubes (1.3 × 16 cm) and incubated at room temperature (20 – 23 C).

Aerobic heterotrophs were enumerated by a 5-tube MPN using 7 ml of R2 medium. The tubes were incubated in the dark and scored for growth by the appearance of visible turbidity up to 4 weeks after inoculation. Fermentative heterotrophs were enumerated by a 5 tube MPN using brain heart infusion in tubes sealed with butyl rubber septa. Positive tubes were scored by turbidity up to 4 weeks after inoculation. SRB were enumerated using 3 tube MPNs in glass tubes sealed with butyl rubber stoppers. Positive tubes were scored by the abundant precipitation of iron sulfides, turbidity, and microscopic checks for growth, as well as the strong odor of sulfide. Some high dilution positive tubes were spot checked for the disappearance of sulfate using a turbidimetric assay (Kolmert et al 2000). FeRB were determined in 3 tube MPNs done in glass vials (15 × 45 mm). The medium contained amorphous Fe-oxides that were produced by the oxidation of FeCl_2_ under microaerobic conditions in a bicarbonate buffered system, pH 6.3 (Neubauer et al 2002). Putative Fe-reduction was determined by measuring the presence of Fe(II) by the ferrozine assay (Stookey 1970), and microscopic evidence for cell growth for up to 2 months. Sulfur-oxidizing bacteria were quantitated in a 3 tube MPN with thiosulfate medium. The presence of S-oxidizing bacteria was determined ( for up to 2 months) by microscopic evidence for growth and in almost all cases by the precipitation of elemental S-particles in the medium. S-disproportionating bacteria were enumerated using a 3 tube MPN series. Putative S-disproportionation was assessed by the formation of a black FeS precipitate, the smell of H_2_S, and microscopic evidence for cell growth for up to 2 months. Neutrophilic Fe-oxidizing bacteria were enumerated using a three tube MPN technique using FeS gradient tubes (Emerson & Moyer 1997). Hydrogen-oxidizing bacteria were assessed in 3 tube MPNs done in Balch tubes using a mineral salts medium (Malik 1981) with a ratio of 80:10:10, N_2_:O_2_:H_2_.

#### Media

Aerobes were grown using R2A medium (Difco Laboratories, #1826-17-1) amended with 2% NaCl and without agar for MPNs. Fermentative bacteria were grown in MPNs on brain heart infusion (Difco Laboratories #0037) with 2% NaCl added, or on plates with Trypticase soy agar with 5% defibrinated sheep’s blood. The composition of the medium for S-oxidizing bacteria (Kuenen et al 1992) was (g/l): NaCl, 25.0 g; (NH_4_)2SO_4_, 1.0 g; MgSO_4_. 7H_2_O, 1.5 g; CaCl_2_, 0.3 g; K_2_HPO_4_, 0.5 g; Na_2_S_2_O_3_. 5H_2_O, 8.0 g; 10 mM HEPES buffer, pH adjusted to 7.2. The medium was filter-sterilized and 1 ml/l of Wolfe’s vitamins and minerals (Wolin et al 1963) stocks solutions were added. The composition of the medium for FeRB (Lovley & Phillips 1988) was: NaCl, 20 g; NaHCO_3_, 2.5 g; MgCl_2_, 2.0 g; CaCl_2_, 0.4 g; KCl, 0.5 g; NH_4_Cl, 0.24 g; KH_2_PO_4_, 0.2; yeast extract, 0.05 g; sodium acetate 0.67 g. The pH of the medium was adjusted to 7.3. It was filter-sterilized and amended with Wolfe’s minerals and approximately 0.1 g per 10 ml of amorphous FeOOH was added. The composition of the medium for S-disproportionaters (Finster et al 1998)was: NaCl, 20g; NaHCO_3_, 2.5g; MgCl_2_ 6H_2_O, 2g; CaCl_2_. 2H_2_O, 0.4g; KCl, 0.5g; NH_4_Cl, 0.25g; KH_2_PO_4_, 0.2g. The pH was adjusted to 7.3, the medium was filter-sterized and stocks of Wolfe’s vitamins and minerals (1 ml/l) were added. After the medium was dispensed (10 ml), approximately 0.1 g of amorphous FeOOH and 0.1 g of flowers of S was added. Following inoculation the tubes were gassed with N_2_:CO_2_ (80:20). The primary medium used for SRB was a lactate medium described in Widdel and Bak (Widdel & Bak 1992) amended with 2.5% NaCl. The medium for *Desulfobacterium* (Widdel & Bak 1992) and related genera was also used with different C-sources as described above.

#### Total cell counts

To determine total cell numbers in the sediments, subsamples of the slurries were fixed with 2% glutaraldehyde. For cell counting, this material was diluted either 1:200 or 1:300 using either sterile phosphate buffered saline or artificial seawater, and 10 *µ*l of this material was smeared evenly within a circle of known diameter on a microscope slide and air dried. A 10 *µ*l solution of 0.25 mM Syto (Molecular Probes, Eugene, OR) was added to the slide and air dried in the dark. Finally 8 *µ*l of sterile PBS was placed on the smear, and a coverslip was added. Using the 100x objective on an Olympus BX 60 microscope 12 – 15 fields were counted per smear (usually between 300 and 700 cells/smear), and three smears per sample were done. The total cell number was determined by taking into account the dilution factors and then normalized to gdw.

## RESULTS

### Total cell numbers

The results of direct cell counts for each of the four sites over 2 1/2 years is shown in Fig 2, and averaged values are shown in Table 2. The total cell number averaged over all the sites at all times was 4.0 × 10^9^ cells. gdw^-1^ sediment. Overall, total cell counts at each of the sites were quite consistent during the course of this study; there were no obvious seasonal fluctuations. The Hog Is. S2 site consistently had the highest cell numbers, and Red Bank consistently had the lowest cell numbers. The Hog Is S1 site, (average = 3.8 × 10^9^ cells; SD = 2.7 × 10^9^) showed the widest fluctuations in total cell number with a range of 1.2 – 9.1 × 10^9^ cells. gdw^-1^. At all sites, most of the cells were associated with sediment particles, and were only visible by microscopy when stained with a DNA-binding fluorescent dye.

**Fig. 2.**
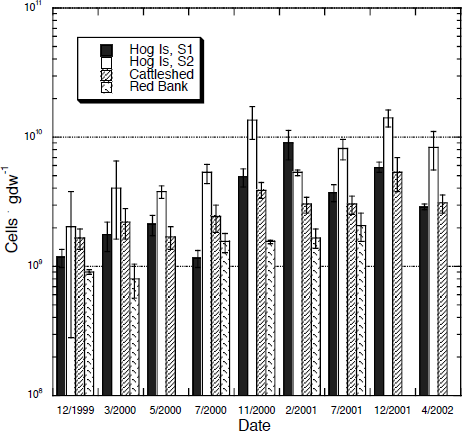
Total direct cell counts for the sampling sites at different times. No counts were done for Red Bank on 5/2000, 12/2001, or 4/2002. The error bars represent the standard deviation.

### MPN studies

The recovery of different populations of bacteria by MPN enumerations are shown in Fig 3 and averaged values in Table2. As is evident there was a wide range in the recovery of different populations between the sites, much greater than the changes documented in total cell numbers. It appeared that the temporal variation within each site was less than the spatial variation between different sites, and there were no obvious seasonal trends in the recoveries of different groups. A synopsis of the results for each of the physiological groups is given below.

**Fig. 3.**
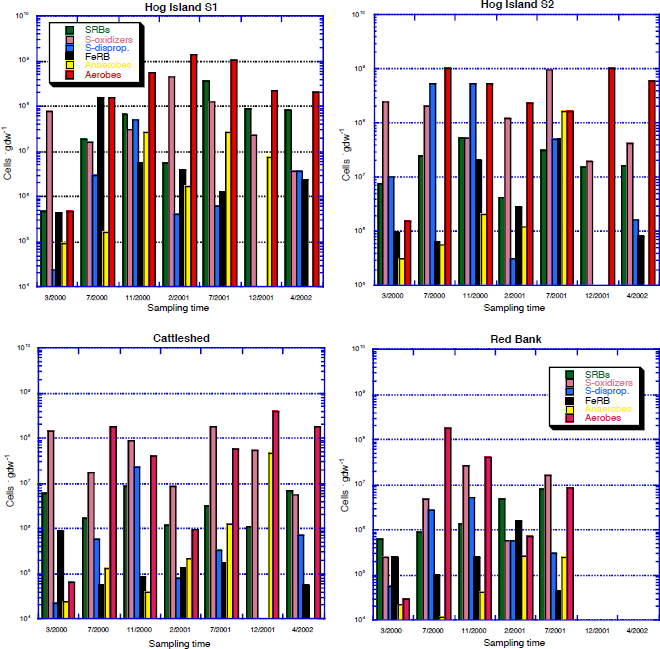
Recoveries of different physiological groups for each site at different times. Numbers of aerobes (3/2000) and anaerobes (3/2000 & 7/2000) were determined by plate counts; on subsequent dates their numbers were determined by MPNs, see text for details. FeRB and S-disproportionaters were not sampled on 12/2001; anaerobe numbers for 4/2002 were lost due to contamination. The Red Bank site was not sampled on 12/2001 and 4/2002.

<u>Aerobic heterotrophs</u>. An important change was made in how aerobes and anaerobes were enumerated during the course of this study. Initially these organisms were quantitated by dilution plating, the results of plating for 3/2000 for aerobes and 3/2000 and 7/2000 for anaerobes are shown in Fig 3. At all subsequent times we used a 5-tube MPN technique and found this gave 100- to 1000-fold increases in total recovered numbers for aerobes, and 10- to 100-fold increases in the numbers of anaerobes compared to plating (Lydell 2002). Thus the numbers used for calculating cell numbers of aerobes and anaerobes are based only on those sampling times when the MPN technique was used. As a result, the aerobic bacteria were the most abundant group recovered overall. The highest numbers were from the Hog Is S1 & S2 sites which had an average of 6 × 10^8^ cells. gdw^-1^, while RB had the lowest numbers.

<u>Sulfur-oxidizers</u>. Thiosulfate-oxidizing bacteria were also abundant with the highest numbers recorded at the Hog Is S2 and S1 sites. Again CS had intermediate numbers and RB yielded the lowest recoveries. The numbers of S-oxidizers were relatively consistent, generally fluctuating by less than an order of magnitude at each site across the seasons. <u>SRB</u>. The highest recoveries of SRB were from the Hog Is. S1 site and the lowest was from Cattleshed and Red Bank, respectively. An initial study on the SRB using different carbon sources, acetate, lactate, butyrate and ethanol indicated that lactate and ethanol yielded the highest recoveries at all these sites. The March, 2000, July 2000, and November 2000 MPN results are combined data from lactate and ethanol MPN’s; however the recoveries on ethanol were always approximately 10-fold lower than those on lactate (results not shown), so later studies were done using lactate only. <u>Sulfur-disproportionating bacteria</u>. This group was present at all the sites at all times of year; however their numbers did fluctuate substantially, note the high SDs in Table 2. For example at the Hog Is S2 site where S-disproportionaters were in the greatest overall abundance (1.6 × 10^8^; SD = 3.1 × 10^8^) their numbers ranged from 5.3 × 10^8^. gdw^-1^ in July and November, 2000 to 3.1 × 10^5^. gdw^-1^ in February, 2001.

<u>Fe-reducers</u>. FeRB were also present at all the sites that were sampled although overall abundances were the lowest of any of the groups sampled. They were most abundant at the Hog Is S1 and S2 sites, and least abundant at Cattleshed, where their numbers were consistently low, 2.4 × 10^5^ cells. gdw^-1^ (SD = 3.4 × 10^5^). Like the S-disproportionating bacteria, their numbers tended to fluctuate more with sampling time than other groups as evidenced by the high standard deviations.

<u>Fermenters</u>. Populations of fermentative anaerobic bacteria were consistently abundant at all the sites, the highest number was recorded in July, 2001 at the Hog Is S2 site, 1.7 × 10^8^ cells. gdw^-1^ (SD = 9.5 × 10^7^), and numbers were generally in the range of 10^6^ - 10^7^ cells. gdw^-1^, except for Red Bank, where numbers were in the 10^5^ range. The numbers shown in Fig 3 for March and July, 2000 were based on plate counts using blood agar; subsequent numbers were based on MPNs using BHI, which gave at least a 10-fold greater recovery from most samples.

<u>Others</u>. MPN quantitation of hydrogen-oxidizing and Fe-oxidizing bacteria indicated these organisms were not abundant members of the community. Fe-oxidizing bacteria were detected at the Red Bank site at numbers of approximately 10^3^ cells. gdw^-1^, but were below this detection limit at the other sites. MPNs for H_2_-oxidizing bacteria showed some growth in the initial dilution series; however transfer of the cultures from high dilutions revealed they did not grow on H_2_; indicating their numbers were below the limits of detection set for the MPNs, which was 10^4^ cells. g^-1^ wet sediment.

### Total recoveries of different metabolic groups

The total recovered cells for all sampling times at the different sites is summarized in Fig 4. At the Hog Is S1 and S2 sites aerobic heterotrophs made up 67% of the combined average population at the S1 site and 55% at the S2 site. Next most abundant at these sites were aerobic S-oxidizing bacteria. At the S1 site these two groups accounted for 76% of the total. At the Cattleshed site these two aerobic groups accounted for 92% of the total recovered populations. The SRB were on average the most abundant of the anaerobic bacteria, accounting for between 2 and 10% of the total recovered populations. At the Hog Is S2 site S-disproportionating bacteria comprised, on average, a larger proportion of the population than SRB, 15% vs. 4%, respectively.

**Fig. 4.**
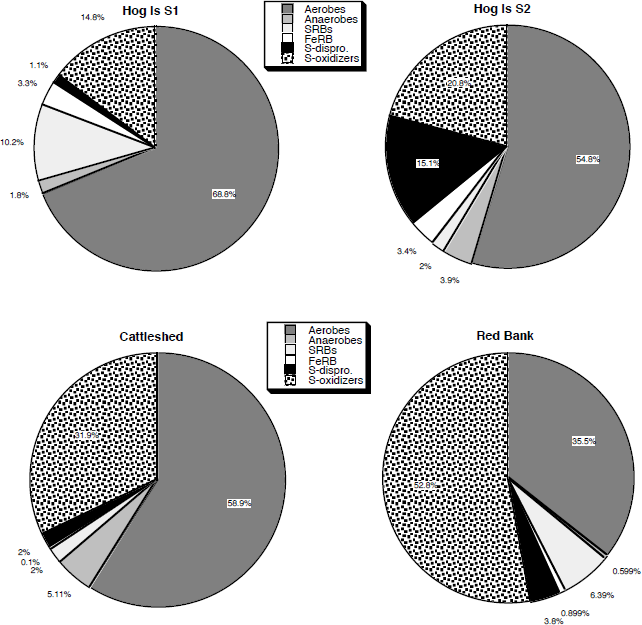
Recovery of different physiological groups at each site as a percentage of the total cultivated population. These numbers are averages of the results shown in Fig 3 and exclude the 12/2001 sampling time.

### Comparison of total cell counts and total recovered cells

Combining all the MPN data at each sampling allowed a comparison of the total number of cells cultured and the total cell direct counts. Two examples of these comparisons, one from winter and one from summer are shown in Fig 5. The numbers and percentages of recovered bacteria at the different sites ranged from a low of 8 × 10^6^ cells. gdw^-1^, representing 0.4% of the total population at Red Bank in July, 2001 to a high of 2 × 10^9^ cells. gdw^-1^ at Hog Is S1 in February, 2001, representing 21% of the total population. The highest percentage recovery for any of the sampling sites was 43% of the total population at Hog Is S1 in July 2001. The two Hog Is sites consistently had the highest numbers of cells recovered averaging about 15% of the total direct count. In contrast, at Red Bank cell recoveries averaged only about 2% or less of the total direct count. At Cattleshed, recoveries were typically between 5 and 10% of the total count.

**Fig. 5.**
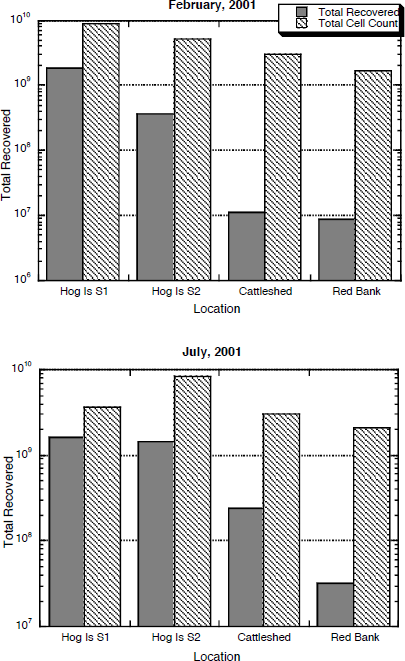
Comparison of total number of bacteria recovered by MPN analysis and the total cell count for the different sites during winter and summer.

## DISCUSSION

### Overall microbial abundance

Total cell numbers at each of the VCR sites were quite constant throughout the 2 years of this study, suggesting that seasonal changes did not dramatically alter total cell numbers. There were significant differences between the total populations at different sites. Total counts ranged from a high of 1.4 × 10^10^ at Hog Is S2 to a low of 7.6 × 10^8^ at Red Bank. In part these differences may be due to differences in the physico-chemical conditions, the Hog Is S2 site had a very high organic content and was not as clay-rich as the Red Bank site. Organic matter alone does not appear to account for all the difference, because at times the Hog Is S1 site had cell numbers nearly as high as S2. Athough S1 was within 3 m of S2, it had a much lower organic C content ( 4% vs 44%), although it did not appear as rich in clay as the Red Bank site.

A study of a salt marsh in North Carolina found cell numbers ranged from a maximum of 1.4 × 10^10^ cells. gdw^-1^ in the top cm of sediment down to about 2 × 10^9^ cells gdw^-1^ at a depth of 10 cm (Rublee & Dornseif 1978). Similar to the results reported here this study also found the cell numbers were quite constant, especially at depth, during a 12 month period. In France, salt marsh sediments taken from a tidal channel were reported to contain a maximum of between 2 and 3 × 10^10^ cells. gdw (Lucas et al 2003) A study of a mid-range salinity (5 –15 ppt) salt marsh with high organic matter on the Hudson River in New York, found total cell numbers of 3 – 4 × 10^10^ cells. gdw^-1^, which are substantially higher numbers than reported here (Burke et al 2002).

### Recoveries of individual metabolic groups

Although salt marsh sediments are thought of as primarily anaerobic habitats, the dominant microbial groups that were recovered during this study were aerobic heterotrophic and thiosulfate-oxidizing bacteria. These results suggest that aerobic metabolism plays an important role in the upper layers of salt marsh sediments. Bioturbation by fiddler crabs and marine invertebrates, and the extensive rhizosphere created by *S. alterniflorans* provide important mechanisms and conduits for transport of both oxygen and labile C into the sediment (Bertness et al 1985, Howes et al 1994). These processes may create a mosaic of oxic habitats ideal for the survival of aerobic and facultative microbes in the top 10 – 20 cm of the salt marsh sediment. It has been estimated that approximately 50% of the O2 taken up by salt marsh sediments is used for aerobic microbial respiration, and the remainder goes toward oxidative processes, primarily S-oxidation (Howes & Teal 1984), where chemical oxidation is supposed to play at least as important a role as biotic oxidation. In part due to methodological considerations, these types of studies based on mass balance calculations have not taken into account microbial population structure, and tend to exclude either bioturbation or rhizosphere transport of O2 as important processes. In light of the high numbers of active aerobic organisms presented here, it may be worth re-evaluating the oxidative respiratory processes in salt marsh sediments, especially vis a vis S-oxidation (see below).

<u>Aerobic heterotrophs</u>. There are few published numbers documenting the abundance of aerobic bacteria in salt marsh sediments. A study of a salt marsh in England dominated by *Spartina townsendii*, found numbers that averaged around 5 × 10^6^ cells. gdw^-1^ (Sivanesan & Manners 1972). At Sapelo Island, Georgia, plate counts typically recovered between 10^5^ and 10^6^ cells. g sediment (wet weight) (Lowe et al 2000). These are similar to the numbers we recovered at the VCR using plate counts. However, as mentioned above, we subsequently found that MPN analysis yielded numbers > 100-fold higher than plate counts, which put recoveries of aerobes in the range of 10^8^ cells. gdw^-1^, with a maximum value of 1.5 × 10^9^ at Hog Is. S1. These latter numbers are in the range of a value of 1.2 × 10^8^ cells. g sediment reported from an MPN enumeration of aerobes from another salt marsh on Sapelo Island, Georgia (Bachoon et al 2001). Preliminary characterization by fatty acid methyl ester analysis indicates that the aerobic heterotrophic bacteria are quite diverse and include representatives from most of the groups of the proteobacteria, the Cytophaga-Flexibacter-Bacteroides (CFB) phylum, as well as gram positive genera. A number of them are likely facultative anaerobes. A more detailed analysis of isolated CFB strains indicates that they are diverse and many appear novel, representing new species and potentially new genera (Lydell et al 2003). Results from clone libraries from DNA extracted directly from the same sediments as were used for MPNs, also show an abundance of presumptive aerobic organisms (Emerson, unpublished results). These results from the VCR sites are consistent with another recent study of a salt marsh in England, where analysis of phospholipid fatty acids (PLFA) profiles during the course of a year found that PFLAs of indicative of aerobic bacteria were consistently more abundant than PFLA’s indicative of SRB (Keith-Roach et al 2002).

<u>Sulfur-oxidizing bacteria</u>. We could find no reliable estimates for numbers of S-oxidizers in salt marsh sediments. The numbers of sulfur-oxidizers at the VCR are among the highest reported for any environment with an overall average number of 1.1 × 10^8^ and an average of 2.3 × 10^8^ at the Hog Is S2 site, which had the highest numbers of S-oxidizers. Imhoff and colleagues found in the range of 10^6^ sulfur-oxidizers gdw^-1^ in shallow marine sediments associated with reed beds in the Baltic Sea (Imhoff et al 1995). Sievert et al found maximal numbers of 1.4 × 10^6^ cells. g wet wt in sediment associated with a shallow hydrothermal vent site (Sievert et al. 1999). Sulfur-oxidizers have also been quantified in rice paddy soil by an MPN technique similar to the one used here, in this case 10^5^ to 10^6^ cells. gdw^-1^ were found (Stubner & Conrad 1998). Based on relative numbers of dominant populations that could be cultivated, our results suggest that S-oxidation is an important process in the marsh. This empirical population data supports mass balance assessments of S-cycling in salt marshes that have previously suggested that chemoautotrophic S-oxidation may contribute substantially to the overall productivity of salt marshes (Howarth 1984, 1993). Almost all of our MPNs were done using thiosulfate as the sole energy source. One comparative MPN series was done using thiosulfate and elemental sulfur. The cell numbers from the elemental sulfur MPNs were 5- to 10-fold lower than those on thiosulfate. Almost all the S-oxidizing bacteria that were enriched through MPNs formed white precipitates of elemental S when grown on thiosulfate, and a number of them accumulated intracellular granules of S. Most are either obligate S-oxidizers, or else grow poorly on heterotrophic media. Analysis of the DNA sequences of the 16S rRNA gene for three of the strains revealed two of them were gamma-proteobacteria not that that closely related to cultivated strains (≤93% similiarity); the third strain was closely related to *Thiobacillus prosperus*.

<u>The sulfur-disproportionating bacteria</u> were present at most sites in numbers of ≥ 10^6^ cells. gdw^-1^, with an overall average of 4.5 × 10^7^ cells. gdw^-1^. At the Hog Is S2 site they were enumerated in maximum numbers of 5.3 × 10^8^ cells. gdw^-1^ in July and November of 2000. To our knowledge there has only been one previous study that quantitated the numbers of S-disproportionaters in both pelagic marine and salt marsh sediments in Denmark (Thamdrup et al 1993). In this case, a maximum of 1.1 × 10^6^ cells/cm^3^ were reported from the salt marsh, taking into account differences in wet wt versus dry wt, these numbers are generally consistent, although on the lower end of the numbers at the VCR. A number of purified enrichments and pure cultures of S-disproportionaters have been obtained from high dilution MPN tubes; however these have not yet been characterized. For example, we do not know what percentage are obligate S-disproportionaters versus being sulfate reducers that are also capable of disproportionation. In any event, these organisms were all enriched under chemolithoautotrophic conditions using elemental S, and given that their numbers are equivalent those of SRB (overall average 3.0 × 10^7^), this provides organismal support to the biogeochemical evidence (Canfield et al 1996) that S-disproportionation is an important pathway for both S‐ and C-cycling in these sediments.

<u>Sulfate reducing bacteria</u>. Literature values for SRB numbers based on MPNs vary widely. One report from Sapelo Island in Georgia found 0.1 to 4 × 10^4^ SRB. g wet wt^-1^, where numbers showed substantial spatial fluctuation both vertically and horizontally (Lowe et al 2000). Another study from Savannah, Georgia found SRB numbers ranged from 10^5^ to 10^6^, which are closer to the numbers we have found at the VCR (Kostka et al 2002b). A comprehensive study by Hines et al enumerated SRB temporally in a marsh in New Hampshire (Hines et al. 1999). Integrating their data for the top 3 cm of bulk salt marsh sediment over time results in about 3.5 × 10^7^ SRB per g wet wt for their study site. Unfortunately, it is not possible to accurately compare these numbers with the numbers from the VCR (overall average = 3.0 × 10^7^. gdw^-1^), since the literature values cited above are based on wet wt of sediment. At the VCR sites, dry wt cell numbers averaged about 3 – 5 times the wet wt numbers. Estimates of SRB from a marsh in Lewes, Delaware using noncultivation-based molecular methods, suggested that averaged total SRB populations could be as high as 1.1 × 10^9^ cells. gdw^-1^ (Edgecomb et al 1999). Using the same technique, based on total DNA extraction, these workers estimated the total bacterial population size in this marsh at 5.2 × 10^10^, which is 3 fold higher than the highest direct counts we have obtained at Hog Island S2. The work of Burke, *et al*, (2002) used FISH probes specific for SRB to enumerate their numbers in a meso-haline marsh and found numbers that averaged 2 – 3 × 10^9^ cells. gdw^-1^, again total direct counts indicated the total population size in this marsh was in the range of 4 – 5 × 10^10^ cells. gdw^-1^. Roony-Varga, et al have found that the dominant SRB that could be cultivated from a marsh in New Hampshire did not appear to be the dominant organisms detected using molecular methods (Rooney-Varga et al 1998). Given the comparisons to molecular results, it is possible we are underestimating the total population sizes of SRB by 5- to 10-fold.

<u>FeRB</u>. Recent work by Kostka, et al, suggests that under conditions where there is both vegetation and extensive bioturbation, Fe-reduction may be as, or more important than sulfate-reduction as a pathway for organic C-mineralization (Kostka et al 2002b). While we were unable to couple process studies with enumeration in the scope of this study, FeRB were abundant and on average were found in comparable numbers to the SRB. The overall average of FeRB at the VCR was 1.7 × 10^7^ cells. gdw^-1^. A study that quantified FeRB in a salt marsh at Sapelo Island using a plating method found on average between 10^5^ and 10^6^ cells/cm^-3^ (Koretsky et al 2003). This is at least circumstantial evidence confirming that Fe-reduction is an important process in the salt marshes and needs further investigation. All MPNs and subsequent enrichments were done using acetate as the growth substrate, physiologically this suggests these organisms are members of the Geobacteracae (Lovley 2000), although we have not as yet characterized the enrichments. <u>Fermentative bacteria</u> were also in the same order of abundance as the other anaerobes when enumerated using the MPN technique, an overall average of 1.2 × 10^7^. While fermentation is generally recognized as an important pathway for C-cycling in salt marshes (King & Wiebe 1980); there are few quantitative analyses of fermenters; most of these have been in the context of investigating the relative contributions of fermenters and SRB to N_2_-fixation (Gandy & Yoch 1988). A study of sediments from an estuary in Scotland reported MPN numbers for fermenters of about 10^3^ cells. gdw^-1^ (Herbert 1975), which is far lower than the numbers reported here; however it is difficult to compare a permanently water-covered estuarine sediment to the more dynamic salt marsh sediment. At the VCR fermentative populations are numerically in balance with anaerobes that respire using organic compounds as electron donors. Based on initial studies approximately two thirds of the isolates obtained under strictly anaerobic conditions were obligate anaerobes. Fatty acid analysis indicates some of the strains are related to *Neisseira* spp, while phenotypic evaluation suggests a number of the strains may also be related to *Clostridium* spp.

### Cultured vs uncultured populations

Over the past decade it has become widely accepted that only 1% or less of the microbes that grow in the environment can be cultured (Pace 1997). However, there are few studies that have explicitly tested this idea by attempting to quantify the important physiological groups from a given environment. The results presented here illustrate the underlying complexity of the problem of ascertaining the culturability of a given community. At the Hog Is sites, which had the greatest total cell numbers and appeared to be most active, the greatest cell recoveries were obtained, typically >10% of the total direct count and as high as 43% of the total count. By contrast, Red Bank, a site with high clay content and little sign of microbial activity, had the lowest total cell counts and total cell recoveries were often <1%. Recognizing the lack of precision in quantifying cell numbers either by MPNs or direct counts in complex samples, these recoveries suggest that a substantial percentage ( >10%) of the population may be culturable using standard techniques at some sites at the VCR.

It should also be kept in mind that a recent study on the culturability of soil bacteria suggested that any dilution-based analysis will under estimate actual numbers by 30-40 % due to the inherent clumping of the cells on particles(Janssen et al 2002). While we have not quantified this for the salt marsh sediments, the same phenomena certainly holds true, since most of the cells are particle associated. It is almost certain that the MPN techniques used here are underestimating individual populations by anywhere between 5 and 10-fold, depending upon site (see discussion of SRB above). However if we assume that this underestimation is consistent across the different physiological groups, then the MPN data should reflect the overall relative populations of different physiological groups within the salt marsh sediment community. Ultimately, it will be the combination of process studies and molecular studies corroborated with cultivation/MPN work that will provide the most efficacy in understanding the functioning of these complex and important ecosystems.

## ACKNOWLEDGEMENTS

We thank Dr. Linda Blum of the University of Virginia for advice and help in accessing the VCR, and the boat captains of the VCR-LTER site for transportation to and from sites under all weather conditions. We are indebted to Dr. Johanna Weiss for the CHN analysis. We thank A. Sit for the gift of elemental sulfur. In addition we thank Dr. Pat Gillevet, Dr. Tom Nerad, Mike Peglar, Jeff Cole, Lori Dowell, and Lindsey Murray for assistance in processing samples. This work was supported in part by NSF grant DEB-9972099.

